## Supplementary figures and tables for "Neo-sex chromosomes, genetic diversity and demographic history in the Critically Endangered Raso lark"

**SUPPLEMENTARY MATERIALS**

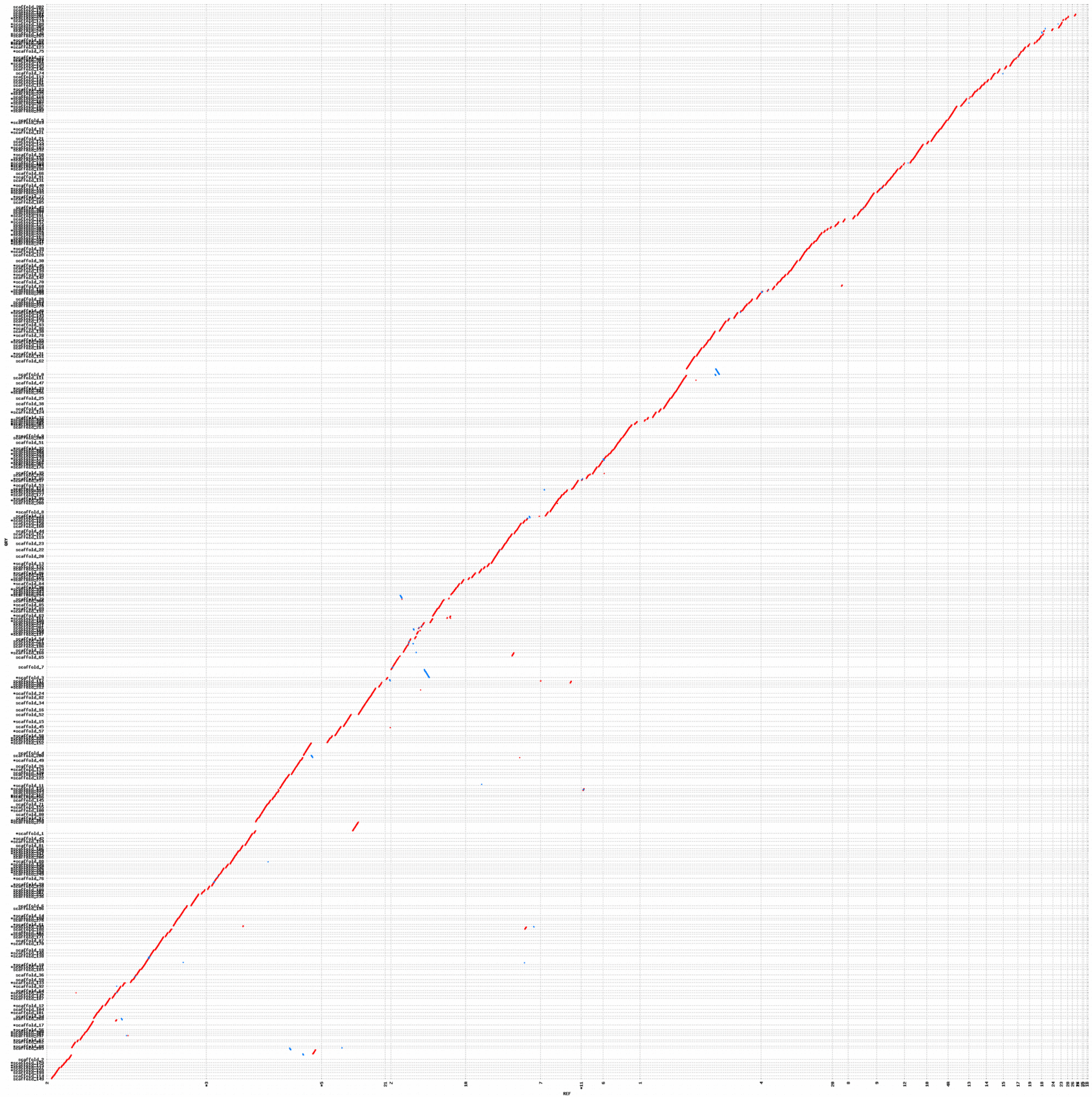

**Figure S1 Alignment between Eurasian skylark scaffolds and zebrafinch chromosomes.** Only scaffolds ≥1 Mb were considered. Points indicate alignments of ≥kb in forward (red) or reverse (blue) orientation. Asterisks indicate scaffolds or chromosomes that were inverted to allow a linear alignment. Most scaffolds map unambiguously to a single chromosome. Exceptions are two of the longest scaffolds, 1 and 2. We conclude that both cases are most likely misassemblies, although this should be confirmed in the future. The final scaffold order used for plots across chromosomes (Figure 2, S2) differed from that shown here by a few manual edits, for example of the location and orientation of scaffold 3.

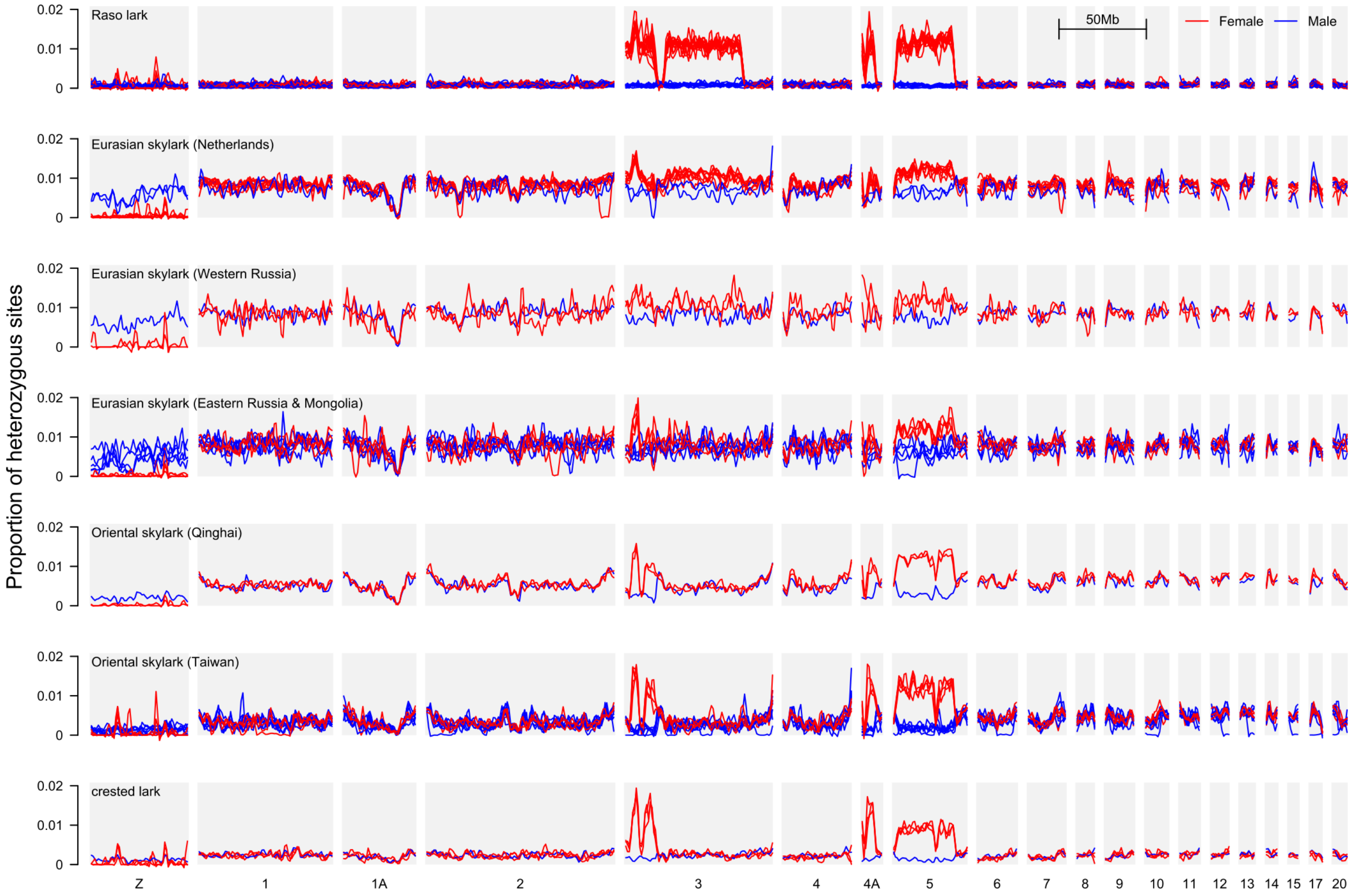


**Figure S2. Individual heterozygosity reveals neo-sex chromosomes.** The proportion of heterozygous sites per 250 kb window in each individual, plotted across each chromosome (based on homology with the zebra finch), with locally-weighted smoothing (loess, span = 10 Mb). Females and males are indicated by red and blue lines, respectively. A lack of heterozygous genotypes in females on the ancestral Z chromosome indicates that it is haploid due to divergence and degradation of the ancestral W (the narrow peaks on the Z most likely reflect collapsed repeats in our assembly). A high density of heterozygous genotypes in females on autosomes is indicative of more recent recombination suppression between neo-W and neo-Z, without significant degradation of the neo-W. Note that the noisier patterns seen in Eurasian skylarks from Western Russia and from Eastern Russia and Mongolia reflect the inclusion of several individuals that only just met the minimum threshold of 3 million sites with ≥5x depth (Table S2).

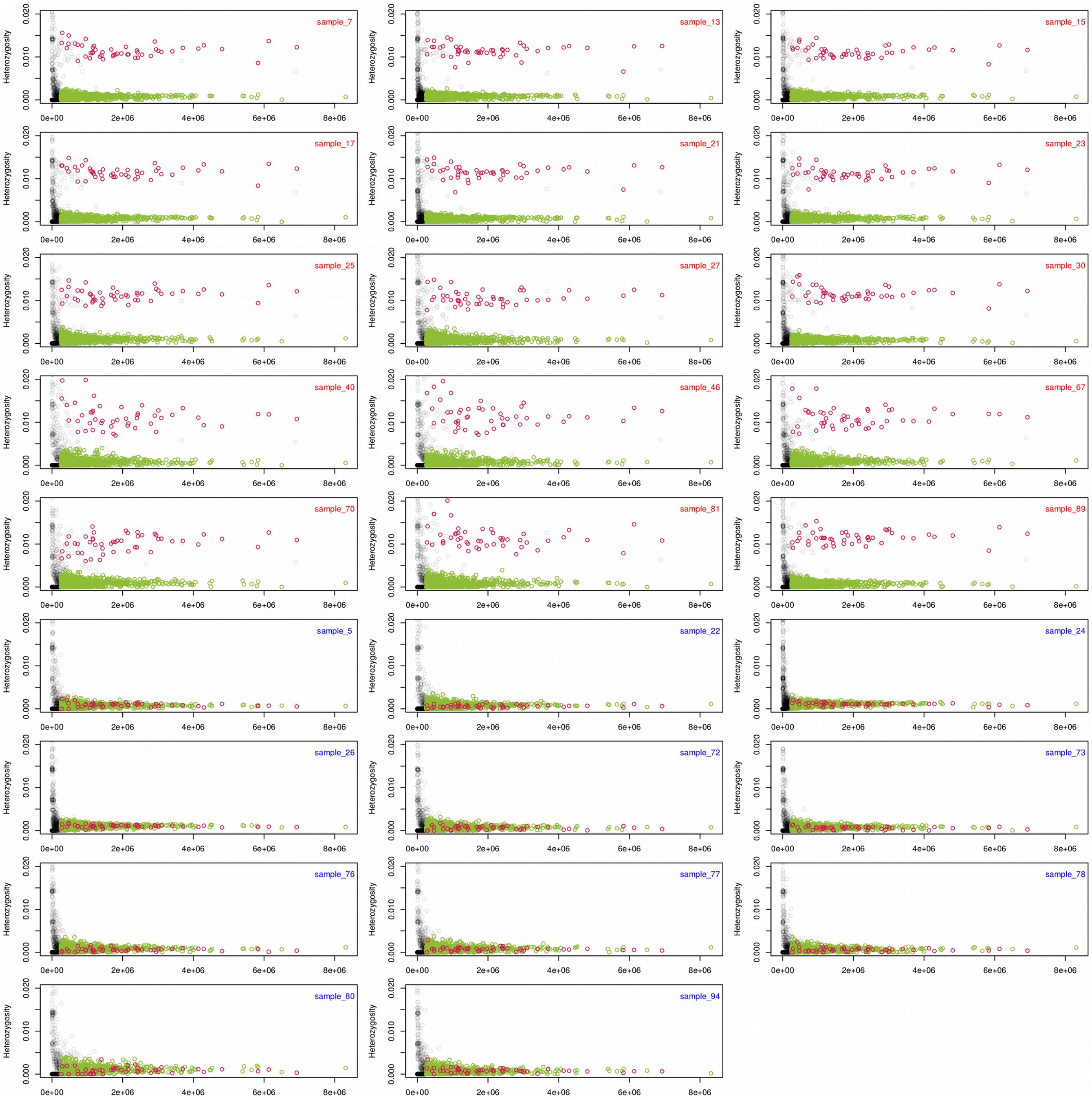


**Figure S3. Identification of scaffolds with suppressed recombination and normal recombination.** Each plot represents a single Raso lark individual, with sample numbers indicated in the top right corner (red=female, blue=male). Points show the proportion of heterozygous genotype calls per scaffold in the given individual (y-axis) plotted against scaffold length (x-axis). We defined normal-recombination scaffolds (green) as those in which the proportion of heterozygous sites is < 0.004 across all samples, and suppressed-recombination scaffolds (magenta) as those in which the proportion of heterozygous sites is > 0.006 in all females and < 0.004 in all males. Scaffolds shorter than 250 kb were excluded.

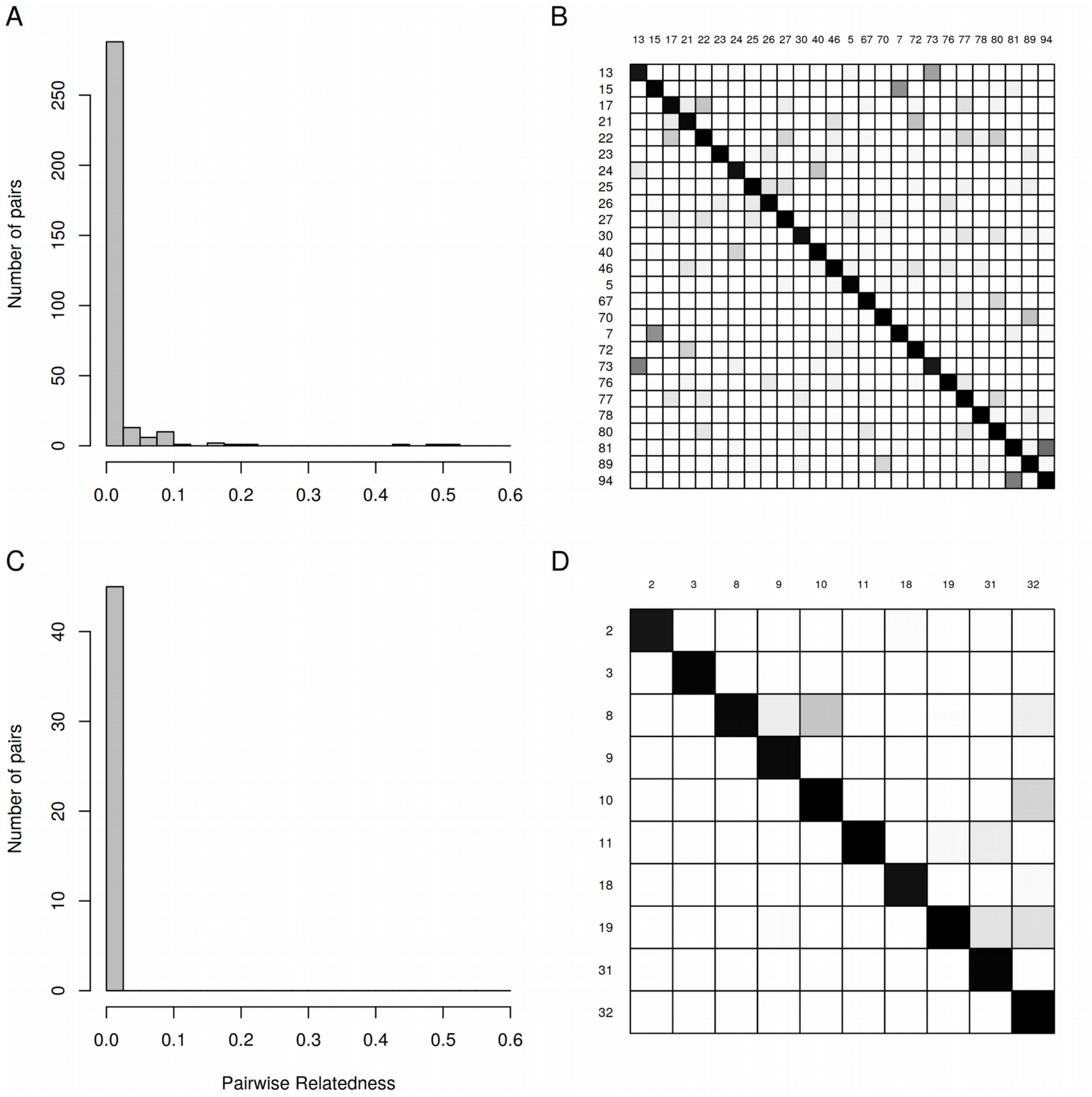


**Figure S4. Relatedness.** (A, C) Histogram of pairwise relatedness ($R_{XY}$) in (A) Raso larks and (C) Eurasian skylarks. (B, D) Pairwise relatedness between all pairs of individuals (lower triangle: $R_{XY}$ [30]; upper triangle and diagonal: G5 [31] in (B) Raso larks and (D) Eurasian skylarks. Shading indicates the level of relatedness, ranging from 0 (unrelated, white) to 1 (completely related, black).

A

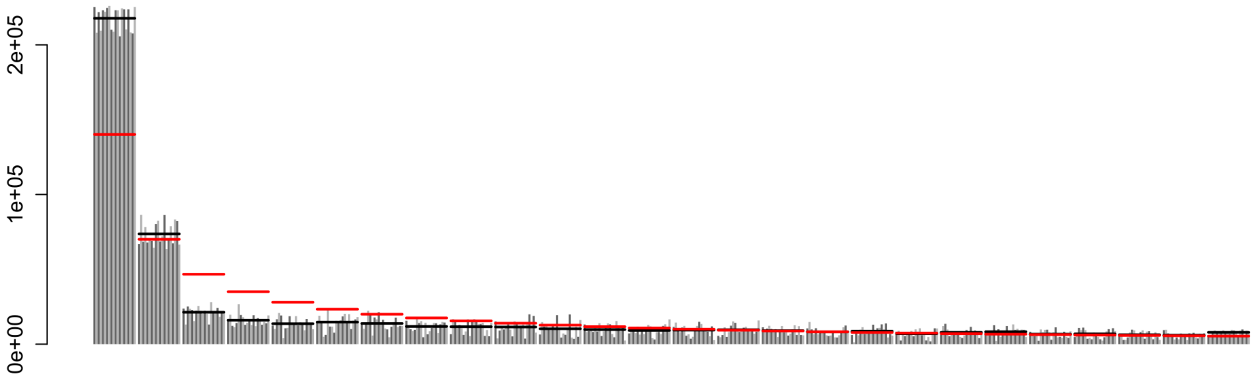


B

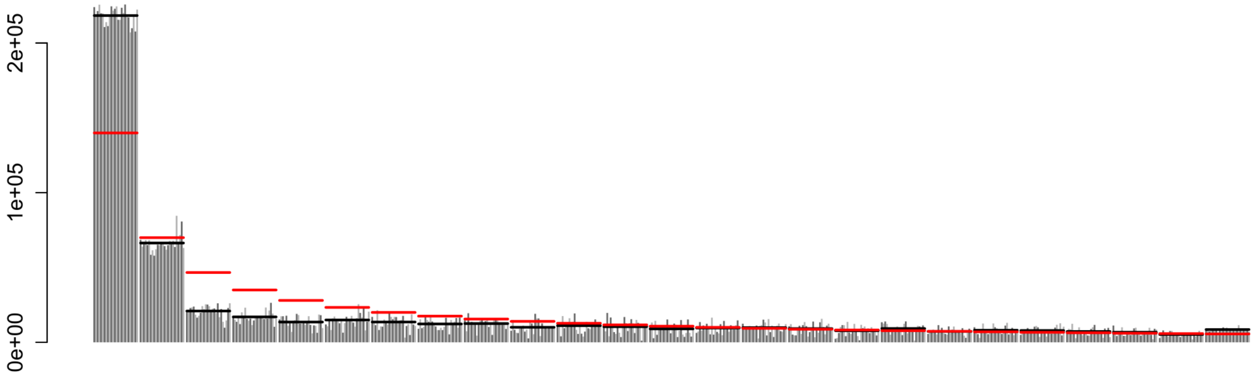


**Figure S5. Consistent excess of rare variants in Raso lark frequency spectra.** A and B both show multiple replicates of the folded site frequency spectrum (SFS) for Raso larks. In A, alternating light and dark shaded bars indicate 20 bootstrap replicates. In B, alternating bars indicate 26 drop-one-out replicates, in which the SFS was recomputed with a single individual excluded in each case. Black horizontal bars indicate the mean across replicates. Red horizontal bars indicate the neutral expectation under constant population size.

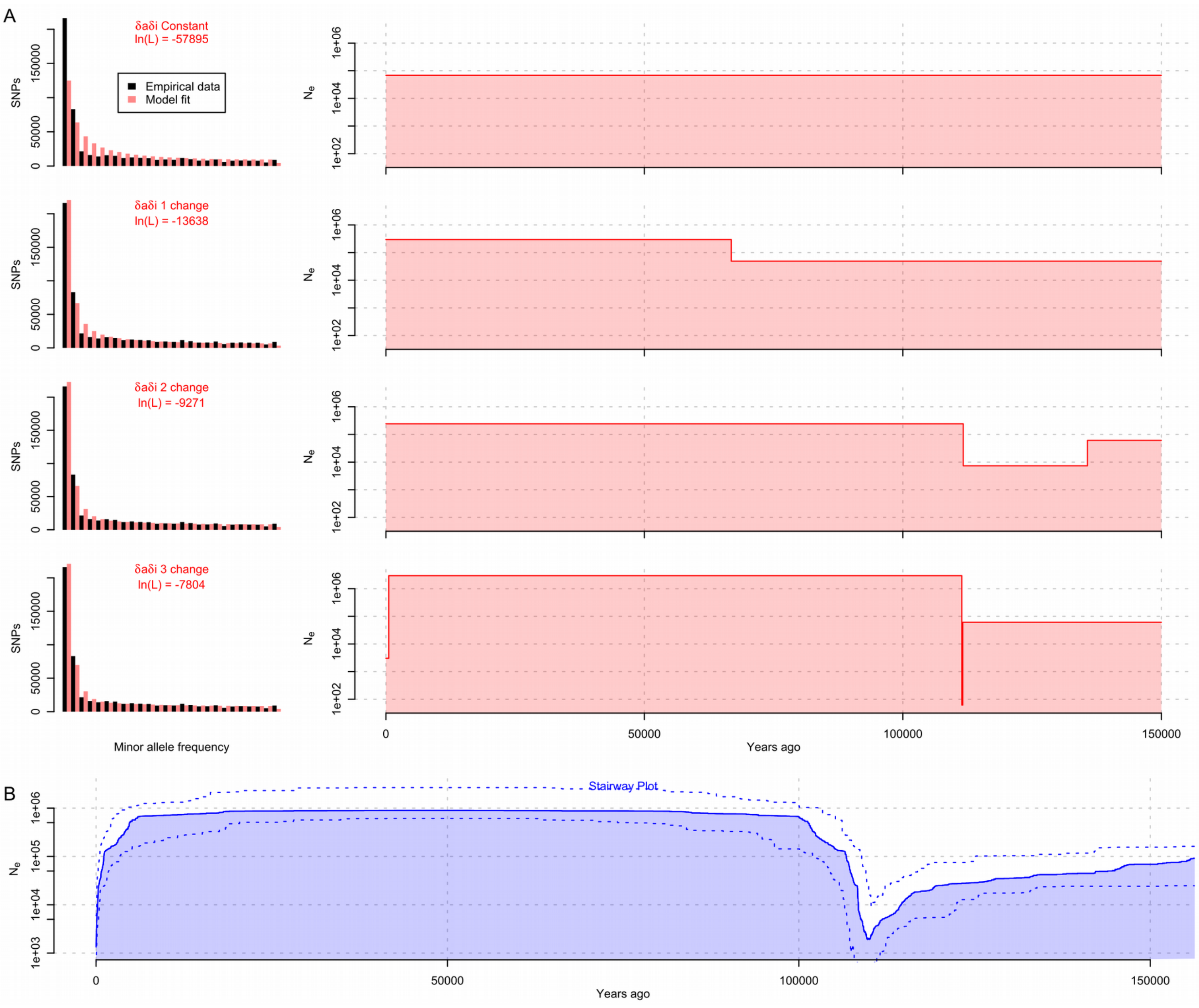


**Figure S6. Demographic model fitting suggests ancient and recent bottleneck.** (A) Four simple demographic models, each allowing a different number of historical changes in population size, were compared using δaδi [33]. Folded allele frequency spectra for the Raso lark RAD-seq data (black) and fitted values under each model (red) are shown on the left. A mismatch between the observed and expected SNP counts indicates a poor fit of the model to the data. The inferred population size is plotted on the right, backwards in time (i.e. the present is on the left). Note that the final contraction in the fourth model is so recent that it cannot be clearly plotted at this scale. (B) Most likely population size history inferred using Stairway Plot [34].

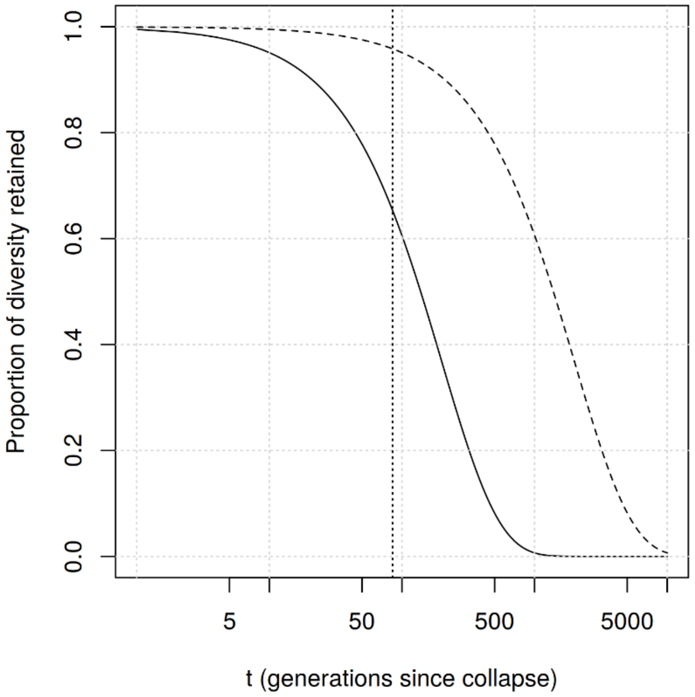


**Figure S7. Expected level of genetic diversity retained after population collapse.** Lines show the expected proportion of pre-existing diversity remaining in a population after a collapse to a $N_e$ of 1000 (dashed line) or 100 (solid line) individuals, after a given number of generations (x axis, log scale). The relative loss of genetic diversity depends only on the current $N_e$, so this plot applies regardless of the historical $N_e$. The vertical dotted line indicates the 85 generation mark assumed for the Raso lark following the1462 human settlement of Cape Verde.

**Table S1. Raso lark population size estimates 1965-2016**

| Year | Population size |
|---|---|
| 1965 | < 50 pairs |
| 1968 | < 40 pairs |
| 1977 | 20 pairs |
| 1981 | 20 pairs |
| 1981 (2nd visit) | 20 pairs |
| 1985 | > 150 pairs |
| 1986* | 200 pairs |
| 1988 | 75-100 birds |
| 1988 (2nd visit) | 250 birds |
| 1989 | 200 birds |
| 1990 | 250 birds |
| 1992 | 250 birds |
| 1998 | 92 birds |
| 2001 | 128-138 birds |
| 2002 | 80-100 birds |
| 2003 | 80 birds |
| 2004 | 57 birds |
| 2005 | 132 birds |
| 2006 | 140 birds |
| 2007 | 159 birds |
| 2008 | 184 birds |
| 2009 | 193 birds |
| 2010 | 486 birds |
| 2011 | 1558 birds |
| 2012 | 1546 birds |
| 2013 | 1314 birds |
| 2014 | 1170 birds |
| 2015 | 900 birds |
| 2016 | 908 birds |
| 2017 | 1514 birds |

Data compiled by references 14 and 16. * indicates first systematic count.

**Table S2. Sample information.**

(Please see separate Excel spreadsheet)

**Table S3. Nucleotide diversity (π) and Watterson's Theta $\Theta_W$ with 95% CI given in brackets**

| | Normal Recombination Regions[1] | | Males only (if possible) | |
|---|---|---|---|---|
| Population | π | $\Theta_W$ | π | $\Theta_W$ |
| Raso lark | 0.001013 (0.001009-0.001018) | 0.001257 (0.001246-0.001266) | 0.000955 (0.000952-0.00096) | 0.001065 (0.001062-0.001069) |
| Eurasian skylark Netherlands | 0.011443 (0.011436-0.011453) | 0.013563 (0.013555-0.013573) | | |
| Eurasian skylark Western Russia | 0.007544 (0.007531-0.007564) | 0.008984 (0.008934-0.009048) | | |
| Eurasian skylark Eastern Russia & Mongolia | 0.007346 (0.007331-0.007355) | 0.008425 (0.008408-0.008439) | 0.006819 (0.006812-0.006827) | 0.007209 (0.007201-0.007217) |
| Oriental skylark Taiwan | 0.004065 (0.004056-0.004074) | 0.004008 (0.003999-0.004015) | 0.00369 (0.003684-0.003699) | 0.003601 (0.003592-0.00361) |

[1] Recombination suppression inferred from heterozygosity in Raso larks (see Figure S3).
